# Development of a novel bioluminescent activity assay for peptide ligases

**DOI:** 10.1101/2021.11.01.466836

**Authors:** Cong-Hui Zhang, Xiao-Xia Shao, Xin-Bo Wang, Li-Li Shou, Ya-Li Liu, Zeng-Guang Xu, Zhan-Yun Guo

**Author notes:** **Correspondence to Z.-Y. Guo**, Research Center for Translational Medicine at East Hospital, School of Life Sciences and Technology, Tongji University, 1239 Siping Road, Shanghai 200092, China.

## Abstract

In recent years, some peptide ligases have been identified, such as bacterial sortases and certain plant asparaginyl or prolyl endopeptidases. Peptide ligases have wide applications in protein labeling and cyclic peptide synthesis. To characterize known peptide ligases or identify new ones, we propose a novel bioluminescent activity assay via the genetic fusion of a recognition motif of a peptide ligase to the C-terminus of an inactive large NanoLuc fragment (LgBiT) and the chemical introduction of a nucleophilic motif preferred by the peptide ligase to the N-terminus of the low-affinity SmBiT complementation tag. When the inactive ligation version LgBiT protein was ligated with the low-affinity ligation version SmBiT tag by the expected peptide ligase, its luciferase activity would be restored and could be quantified sensitively according to the measured bioluminescence. In the present study, we first validated the novel bioluminescent activity assay using bacterial sortase A and plant butelase-1. Subsequently, we screened novel peptide ligases from crude extracts of selected plants using two LgBiT–SmBiT ligation pairs. Among 80 common higher plants, we identified that five of them likely express asparaginyl endopeptidase-type peptide ligase and four of them likely express prolyl endopeptidase-type peptide ligase, suggesting that peptide ligases are not so rare in higher plants and more of them await discovery. The novel bioluminescent activity assay is ultrasensitive, convenient for use, and resistant to protease interference, and thus would have wide applications for characterizing known peptide ligases or screening new ones from various sources in future studies.

## 1. Introduction

Natural peptide ligases (or protein ligases) are a group of special endoproteases, such as the bacterial sortases [1–4] and certain plant asparaginyl endopeptidases (AEPs, also known as peptidyl asparaginyl ligases, PALs) [5–7] and prolyl endopeptidases (PEPs, also known as prolyl oligopeptidase, POP) [8–12], that catalyze polypeptide chain ligation via a transpeptidation mechanism. Sortases are present widely in gram-positive bacteria, and are responsible for the covalent attachment of specific cell surface proteins to the cell wall [1–4]. Sortases are required for the pathogenesis of some pathogens, and thus represent promising drug targets to treat their related infections [13–15]. Sortase A from *Staphylococcus aureus* is the prototype sortase, and was first identified in 1999 [16]. Certain plant AEPs, such as butelase-1 from *Clitoria ternatea* (butterfly pea) [17] and OaAEP1 from *Oldenlandia affinis* [18], and certain plant PEPs, such as PCY1 from *Saponaria vaccaria* [8] and POPB from *Galerina marginata* [11], were also identified as natural peptide ligases recently. They catalyze peptide-chain cyclization and are responsible for the biosynthesis of certain cyclic peptides from genetically encoded linear peptide precursors [5–12].

Peptide ligases have been widely used in protein labeling and cyclic peptide synthesis in recent years [19–25]. To characterize known peptide ligases or identify new ones, in the present study we developed a novel bioluminescent activity assay based on an inactive large NanoLuc fragment (LgBiT) and a low-affinity SmBiT complementation tag. The low-affinity LgBiT–SmBiT pair was recently developed by Promega to monitor dynamic protein–protein interactions and was termed NanoLuc Binary Technology (NanoBiT) [26]. When the inactive LgBiT protein is complemented with the low-affinity SmBiT tag or a high-affinity HiBiT tag, it restores high luciferase activity [26], resulting in a detection limit of femtomoles, or even attomoles, on most microplate readers measured using 96-well or 384-well plates.

The free SmBiT tag can only efficiently complement with the LgBiT protein at high concentrations because of its low binding affinity with LgBiT (K_d_ = 190 μM) [26]. We hypothesized that once the low-affinity SmBiT tag was covalently attached to the C-terminus of the inactive LgBiT protein, it would also efficiently restore the luciferase activity of LgBiT. Moreover, activity of the SmBiT-attached LgBiT would be no longer dependent upon the free SmBiT. Based on this hypothesis, we proposed a novel bioluminescent activity assay that might be suitable to assay various peptide ligases. To develop this bioluminescent activity assay for one or a group of peptide ligases, a known or deduced recognition motif of the peptide ligase(s) was genetically fused to the C-terminus of the inactive LgBiT protein (Fig. 1A), and a known or deduced nucleophilic motif preferred by the peptide ligase(s) was chemically introduced to the N-terminus of the synthetic SmBiT tag (Fig. 1B). Once the inactive ligation version LgBiT protein was ligated with the low-affinity ligation version SmBiT tag by the expected peptide ligase(s), its luciferase activity would be restored and could be quantified accurately according to the measured bioluminescence (Fig. 1C). As a result, ligation activity of the peptide ligase(s) would be conveniently monitored according to bioluminescence of its ligation product.

**Fig. 1.**
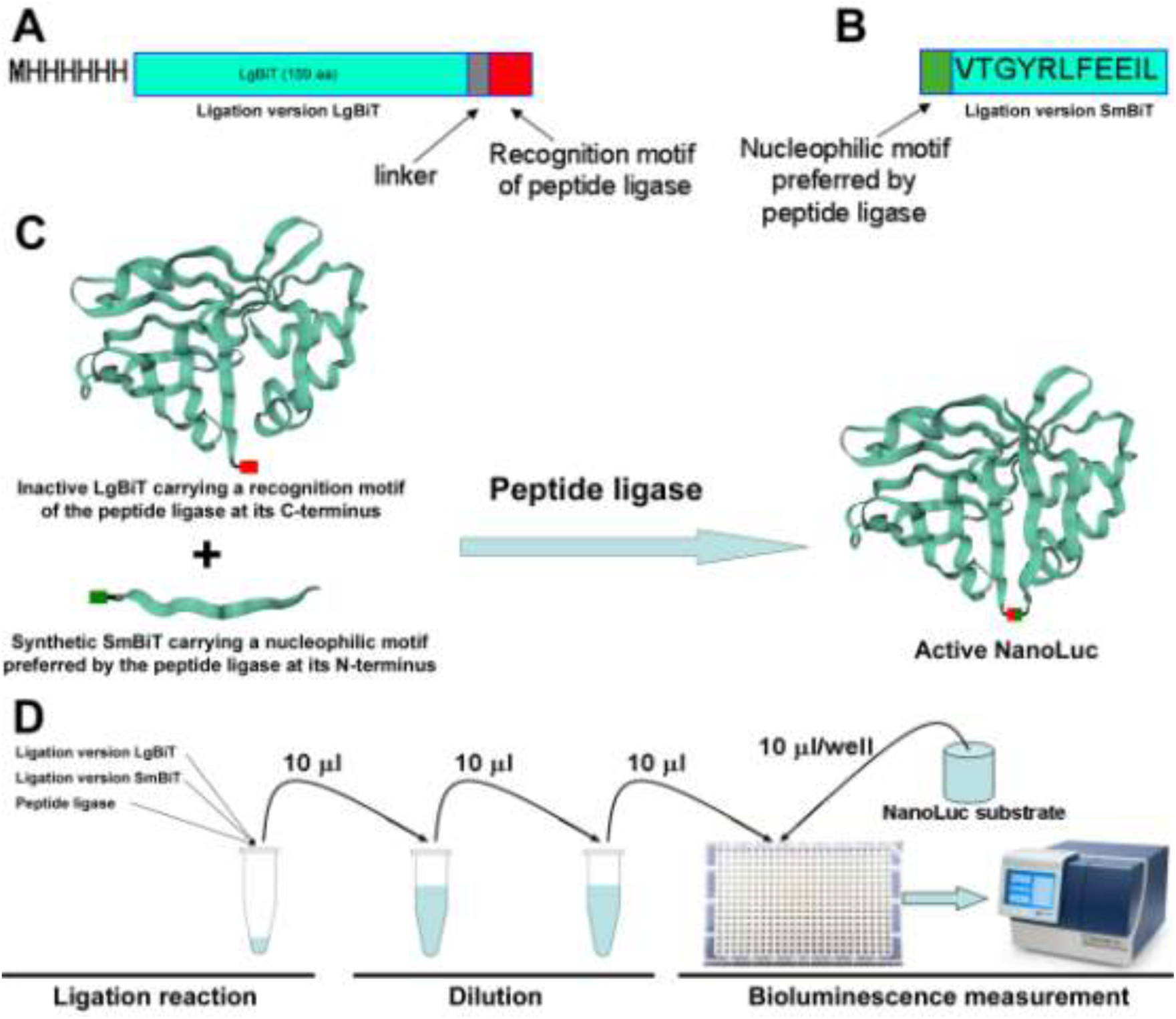
Proposal of a novel bioluminescent activity assay for peptide ligases. (A) Schematic presentation of the ligation version LgBiT protein carrying a known or deduced recognition motif of a peptide ligase at its C-terminus. (B) Schematic presentation of the ligation version SmBiT tag carrying a known or deduced nucleophilic motif preferred by the peptide ligase at its N-terminus. Amino acids in LgBiT and SmBiT are shown using the one-letter code. (C) Schematic presentation of the peptide ligase-catalyzed ligation of the ligation version LgBiT and the ligation version SmBiT. The structural model of LgBiT was adopted from the crystal structure of NanoLuc (PDB ID: 5IBO). (D) Flowchart showing the steps involved in the novel bioluminescent peptide ligase activity assay.

In general, peptide ligases require high substrate concentration (typically micromolar range) for efficient catalysis of the ligation reaction (Fig. 1D). However, micromolar concentrations of LgBiT and SmBiT would form significant intermolecular complementation and produce strong background bioluminescence that would be much higher than the signal bioluminescence produced by the ligation product (Table S1). To solve this problem, we introduced an essential dilution step between the ligation reaction and the bioluminescence measurement (Fig. 1D), because dilution drastically lowers the background bioluminescence (Table S1). After appropriate dilution, the peptide ligase activity can be reliably detected even when only 0.1% of the LgBiT substrate is converted to the ligation product (Table S1).

In the present study, we first validated the novel bioluminescent activity assay using the bacterial sortase A and the plant butelase-1, and then screened novel peptide ligases from crude extracts of 80 common higher plants, either using LgBiT-NHV and GI-SmBiT as a ligation pair or using LgBiT-PIQ and FG-SmBiT as a ligation pair. We identified possible AEP-type peptide ligase activity in five evolutionarily distant plants and possible PEP-type peptide ligase activity in four evolutionarily distant plants, implying that peptide ligases are not so rare in higher plants, and more of them are waiting to be discovered in future studies via an efficient screening approach. The present study demonstrated that the novel bioluminescent activity assay is ultrasensitive, convenient for use, and resistant to interference of proteases, thus it would have wide applications for characterizing known peptide ligases or screening new ones from crude extracts of plants or other sources in future studies.

## 2. Materials and Methods

### 2.1. Chemical synthesis of the complementation tags

The ligation version low-affinity SmBiT tags (4G-SmBiT, GI-SmBiT, and FG-SmBiT) and the high-affinity HiBiT tag were chemically synthesized at GL Biotech (Shanghai, China) via solid-phase peptide synthesis using standard Fmoc methodology. The synthetic peptides with an amidated C-terminus were purified to homogeneity using high performance liquid chromatography (HPLC) with a semi-preparative C_18_ reverse-phase column (Zorbax 300SB-C18, 9.4 × 250 mm; Agilent Technologies, Santa Clara, CA, USA), and confirmed by electrospray mass spectrometry. The purified complementation tags were dissolved in 1.0 mM aqueous hydrochloride (pH 3.0) and quantified by their UV absorbance at 280 nm using the extinction coefficient of 1490 M^-1^cm^-1^ for the ligation version SmBiT tags and 5500 M^-1^cm^-1^ for the HiBiT tag. The peptide stock solution was then aliquoted (20–50 μl/tube) and stored at –80°C for later activity assays.

### 2.2. Overexpression and purification of the LgBiT proteins

The nucleotide sequence of the original LgBiT protein was chemically synthesized at GeneWiz (Souzhou, China) according to its published sequence [26]. After cleavage with restriction enzymes NdeI and EcoRI, the synthetic DNA fragment was ligated into a pET expression vector, resulting in the construct pET/LgBiT, which encodes an N-terminally 6×His-tagged LgBiT protein (Fig. S1). The nucleotide sequences of the ligation version LgBiT proteins and the expected ligation products were generated by PCR amplification of the LgBiT sequence and subsequent cloning into the pET vector. The LgBiT coding sequence in all expression constructs was confirmed by DNA sequencing.

Thereafter, the LgBiT proteins were overexpressed in *Escherichia coli* strain BL21(DE3) according to standard protocols. Briefly, the transformed *E. coli* cells were cultured in liquid Luria-Bertani (LB) medium (plus 100 μg/ml ampicillin) to OD_600_ = 1–2 at 37 °C with vigorous shaking (250 rpm). Thereafter, isopropyl β-D-thiogalactoside was added to a final concentration of 0.5 mM, and the cells were continuously cultured at 16 °C for 12–16 h with gentle shaking (100 rpm). Subsequently, the cells were harvested by centrifugation (5000 *g*, 10 min), resuspended in lysis buffer (20 mM phosphate buffer, pH 7.4, 0.5 M NaCl), and lysed by sonication. After centrifugation (12000 *g*, 30 min), the supernatant was applied to an immobilized metal ion affinity chromatography, and the LgBiT protein was eluted from the Ni^2+^ column using 250 mM imidazole (in lysis buffer). Thereafter, the eluted LgBiT fraction was dialyzed against 10 mM Tris–Cl buffer (pH 7.5), and then applied to an ion-exchange chromatography. The LgBiT protein was eluted from the DEAE-Sepharose Fast Flow column using a linear NaCl gradient (0–0.2 M NaCl in 10 mM Tris–Cl buffer, pH 7.5). The eluted LgBiT fraction was analyzed by sodium dodecyl sulfate-polyacrylamide gel electrophoresis (SDS-PAGE) and quantified by its UV absorbance at 280 nm using an extinction coefficient of 20000 M^-1^cm^-1^. The purified LgBiT protein was then aliquoted (20–50 μl/tube) and stored at –80 °C for later activity assays.

### 2.3. Luciferase activity measurement of the LgBiT proteins

For the intrinsic luciferase activity assay, the purified LgBiT protein was diluted sequentially using the diluting buffer [phosphate-buffered saline (PBS) plus 0.1% bovine serum albumin (BSA) and 0.01% Tween-20] to 10 nM, 1.0 nM, and 0.1 nM, respectively. Thereafter, 10 μl of the diluted samples were transferred to a 384-well plate. For the HiBiT complementation assay, the purified LgBiT protein was diluted sequentially using the diluting buffer to 20 nM, 2.0 nM, and 0.2 nM, respectively. Thereafter, the diluted samples were mixed with an equal volume of 80 nM of synthetic HiBiT (in diluting buffer). After incubation at room temperature for ∼10 min, 10 μl of the mixture was transferred to a 384-well plate. Finally, freshly diluted NanoLuc substrate (100-fold dilution using the diluting buffer) was added (10 μl/well), and bioluminescence was immediately measured on a SpectraMax iD3 microplate reader (Molecular Devices, Sunnyvale, CA, USA) under the luminescence mode. The measured bioluminescence data are expressed as the mean ± standard deviation (SD, *n* = 3) and plotted using the SigmaPlot10.0 software (Systat Software, Chicago, IL, USA).

### 2.4. Overexpression and purification of the circular sortase A

The circular sortase A was prepared according to our previous report [27]. Cyclization of the N-terminally truncated *Staphylococcus aureus* sortase A (residues 60–206) was mediated by a split mini-intern derived from *Synechocystis* sp. PCC6803 DnaB according to a previous study [28]. The circular sortase A was overexpressed in *E. coli* as a soluble protein and then purified by sequential DEAE anion ion-exchange chromatography and SP cation ion-exchange chromatography, according to our previous study [27]. The purified circular sortase A was analyzed by SDS-PAGE and quantified by UV absorbance at 280 nm using an extinction coefficient of 13490 M^-1^cm^-1^. The circular sortase A fraction was then aliquoted (20–50 μl/tube) and stored at –80 °C for later ligation assays.

### 2.5. Preparation of the crude extract from butterfly peas and other plants

Butterfly peas (*Clitoria ternatea*) were cultivated in Hainan province of China, and the tender leaves and stems were collected, transported, and stored at –80 °C. Other higher plants were grown around the campus of Tongji University, and their leaves and/or flowers were collected and used as fresh. To prepare the crude extract from butterfly peas, 1.0 g of the frozen leaves and stems was ground in a cold mortar with 4.0 ml of cold PBS or the previously reported reductive buffer (20 mM phosphate, pH 6.5, 1.0 mM EDTA, and 5.0 mM β-mercaptoethanol) [17,29]. To prepare the crude extract from other plants, 1.0 g of their fresh leaves and/or flowers was ground in a cold mortar with 2.0 ml of cold PBS. After centrifugation (10000 *g*, 5 min), the supernatant was used as crude extract for the ligation activity assay.

### 2.6. Partial purification of butelase-1 from butterfly peas

Partial purification of butelase-1 from the leaves and stems of butterfly peas was conducted according to a previously published procedure [17,29]. The final partially purified product was analyzed by SDS-PAGE and quantified according to its band density using BSA as a standard. The partially purified butelase-1 fraction was then aliquoted (20–50 μl/tube) and stored at –80 °C for ligation assays.

### 2.7. Peptide ligase activity assays

To conduct the sortase A activity assay, the purified LgBiT-long-LPETG (eluted from the DEAE column), the synthetic 4G-SmBiT (dissolved in 1.0 mM aqueous HCl), and the purified circular sortase A (eluted from the SP column) were sequentially mixed together at the indicated final concentrations in the sortase A ligation buffer (100 mM Tris–Cl, pH 7.5, 10 mM CaCl_2_, with or without 0.1% BSA, as indicated). To conduct the butelase-1 activity assay, the purified LgBiT-NHV (eluted from the DEAE column), the synthetic GI-SmBiT (dissolved in 1.0 mM aqueous HCl), and the crude butterfly pea extract or partially purified butelase-1 were sequentially mixed together at the indicated final concentrations in PBS buffer (PBS with or without 0.1% BSA, as indicated) or the reductive buffer (20 mM phosphate, pH 6.5, 1.0 mM EDTA, and 5.0 mM β-mercaptoethanol). To screen the peptide ligase activity from the crude extracts of other plants, the ligation pair of LgBiT-NHV and GI-SmBiT, or the ligation pair of LgBiT-PIQ and FG-SmBiT, was mixed with the crude extract at the indicated final concentrations in PBS. The ligation reaction was conducted at 25°C. At the indicated times, 10 μl of the reaction mixture was removed and diluted into 1.0 ml of the diluting solution (PBS plus 0.1% BSA and 0.01% Tween-20). If necessary, further dilution (10–1,000-fold) was conducted. Finally, the diluted samples were transferred to a 384-well plate (10 μl/well), and their bioluminescence was measured immediately on a SpectraMax iD3 microplate reader after addition of the freshly diluted NanoLuc substrate (100-fold dilution using the diluting buffer, 10 μl/well). According to the measured bioluminescence and specific activity of the ligation product, the concentrations of the ligation product in the ligation mixture were calculated. The measured data are expressed as the mean ± SD (*n* = 3) and plotted using the SigmaPlot10.0 software (Systat Software).

## 3. Results

### 3.1. Validation of the novel bioluminescent activity assay using bacterial sortase A

To validate the bioluminescent activity assay using bacterial sortase A, we first introduced a sortase A-recognition motif (LPETG) into the C-terminus of LgBiT via a short peptide linker (Fig. 2A). Unfortunately, the resultant LgBiT-LPETG displayed high intrinsic luciferase activity (approximately 30% of that complemented with HiBiT), implying this ligation version is unsuitable for the peptide ligase activity assay. To eliminate its intrinsic luciferase activity, we introduced a longer flexible linker comprising 18 amino acids, and the resultant ligation version was designated as LgBiT-long-LPETG (Fig. 2A). After overexpression and purification, this ligation version LgBiT protein displayed a major band with the expected molecular weight as analyzed by SDS-PAGE (Fig. 2B, inner panel). In the luciferase activity assay, LgBiT-long-LPETG emitted strong bioluminescence after complementation with the synthetic HiBiT tag, with a linear range of several orders of magnitude (Fig. 2B). Its calculated specific activity reached 5.5 × 10^5^ RLU/fmol when measured on an iD3 microplate reader using a 384-well plate, which was almost identical to that of the native LgBiT protein (Fig. 2B). Without HiBiT complementation, its bioluminescence was ∼1000-fold lower (Fig. 2B). Thus, LgBiT-long-LPETG was a suitable version for developing the bioluminescent activity assay for sortase A.

**Fig. 2.**
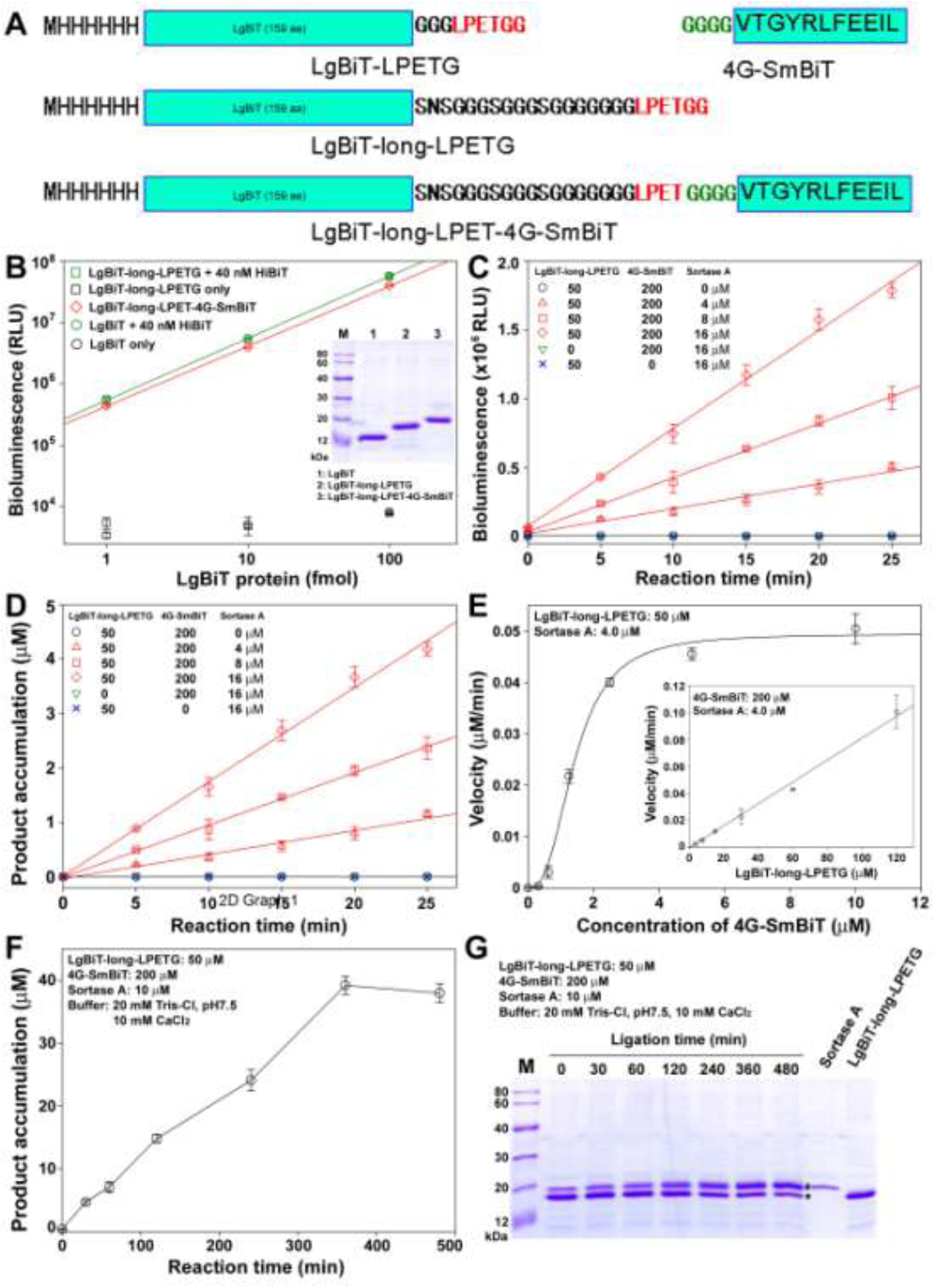
Validation of the bioluminescent activity assay using bacterial sortase A. (A) Schematic presentation of the ligation pair for the sortase A activity assay. Amino acids in these molecules are shown using the one-letter code. (B) Luciferase activity measurement of the recombinant LgBiT proteins. Inner panel, SDS-PAGE analysis. Samples (2 μg/lane) were loaded onto a 15% SDS-PAGE gel, and the gel was stained using Coomassie brilliant blue R250 after electrophoresis. (C) Monitoring the sortase A-catalyzed ligation using bioluminescence. The reaction components at the indicated concentrations were mixed together in the reaction buffer with 0.1% BSA. At the indicated times, 10 μl of the reaction mixture was removed and diluted 10,000-fold using the diluting buffer. Thereafter, bioluminescence of diluted samples was measured on an iD3 microplate reader using a 384-well plate after addition of the diluted NanoLuc substrate (10 μl/well). The measured bioluminescence data are expressed as the mean ± SD (*n* = 3) and fitted to linear curves using the SigmaPlot10.0 software. (D) Ligation product accumulation in the sortase A-catalyzed ligation. The bioluminescence data in panel (C) were converted to ligation product concentration according to the specific activity of the recombinant ligation product. (E) Determination of the K_m_ value of the circular sortase A towards 4G-SmBiT. Inner panel: Determination of the K_m_ value of the circular sortase A towards LgBiT-long-LPETG. The ligation reaction was conducted in the reaction buffer with 0.1% BSA, and the initial velocity was calculated from the ligation product generated within 5 min according to the measured bioluminescence and the specific activity of the ligation product. The measured data for 4G-SmBiT were fitted to the function of V = V_max_[S]^n^/(K_m_ ^n^+[S]^n^), and those for LgBiT-LPETG were fitted to a linear curve, using SigmaPlot10.0 software. (F,G) Monitoring the sortase A-catalyzed ligation by both bioluminescence (F) and SDS-PAGE (G). The ligation reaction was conducted in the reaction buffer without BSA. At the indicated times, 10 μl of the reaction mixture was removed and either diluted 100,000-fold using the diluting buffer for bioluminescence measurement or mixed with 40 μl of the loading buffer for SDS-PAGE analysis. Thereafter, 10 μl of the diluted sample was used for the bioluminescence measurement or for SDS-PAGE analysis.

To develop the bioluminescent activity assay for sortase A, a prerequisite is that the ligation product of LgBiT-long-LPETG and 4G-SmBiT acquired high luciferase activity. To test this hypothesis, we overexpressed and purified the expected ligation product, designated as LgBiT-long-LPET-4G-SmBiT (Fig. 2A). The luciferase activity assay demonstrated that the recombinant ligation product was highly active (Fig. 2B), with a calculated specific activity only slightly lower than that of the HiBiT-complemented LgBiT-long-LPETG (4.1 × 10^5^ RLU/fmol versus 5.5 × 10^5^ RLU/fmol).

Thereafter, we tested whether LgBiT-long-LPETG and 4G-SmBiT could be ligated together by sortase A. After the recombinant LgBiT-long-LPETG, the synthetic 4G-SmBiT, and the recombinant circular sortase A were mixed together, the measured bioluminescence increased in a linear manner within 25 min (Fig. 2C). Moreover, the bioluminescence increase rate was proportional to the sortase A concentration (Fig. 2C). In contrast, no bioluminescence increase was observed in the absence of the circular sortase A, or LgBiT-long-LPETG, or 4G-SmBiT (Fig. 2C). Thus, it seemed that the observed bioluminescence increase was caused by sortase A-catalyzed ligation of the recombinant LgBiT-long-LPETG and the synthetic 4G-SmBiT. According to the measured specific activity of the ligation product, the bioluminescence data were converted to ligation product concentration (Fig. 3D). Under these reaction conditions, micromolar amounts of the ligation product were produced within 25 min, with a calculated catalysis efficiency of approximately 0.01 min^-1^, similar to the reported k_cat_ value of the circular sortase A measured using short peptide substrates [28]. Thus, it seemed that sortase A can efficiently utilize LgBiT-long-LPETG and 4G-SmBiT as substrates.

**Fig. 3.**
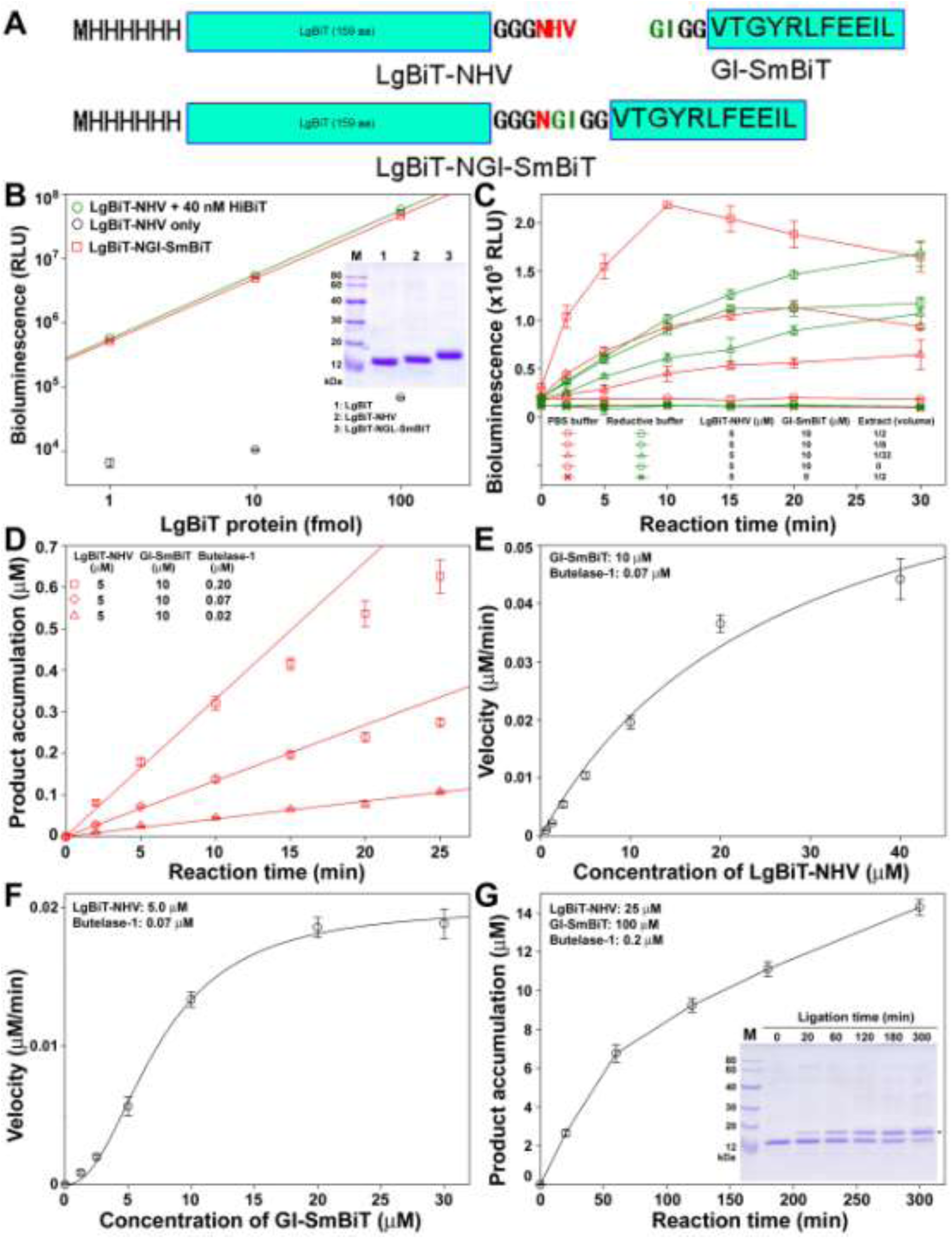
Validation of the bioluminescent activity assay using plant butelase-1. (A) Schematic presentation of the ligation pair for the butelase-1 activity assay. Amino acids in these molecules are shown using the one-letter code. (B) Luciferase activity measurement of the recombinant LgBiT proteins. Inner panel: SDS-PAGE analysis. Samples (2 μg/lane) were loaded onto a 15% SDS-PAGE gel, and the gel was stained using Coomassie brilliant blue R250 after electrophoresis. (C) Bioluminescence change in the ligation of LgBiT-NHV with GI-SmBiT catalyzed by the crude extract of butterfly peas. The ligation reaction was conducted either in PBS or in reductive buffer without BSA. At the indicated times, 10 μl of the reaction mixture was removed and diluted 3,000-fold using the diluting buffer. Thereafter, bioluminescence of the diluted samples (10 μl/well) was measured on an iD3 microplate reader using a 384-well plate after addition of the diluted NanoLuc substrate (10 μl/well). The measured bioluminescence data are expressed as the mean ± SD (*n* = 3) and plotted using the SigmaPlot10.0 software. (D) Ligation product accumulation catalyzed by partially purified butelase-1. The ligation reaction was conducted in PBS with 0.1% BSA. At the indicated times, 10 μl of the reaction mixture was removed and diluted 10,000-fold using the diluting buffer. Concentrations of the ligation product in the ligation mixture were calculated from the measured bioluminescence data according to specific activity of the recombinant ligation product. (E,F) Enzymatic kinetics of butelase-1 towards LgBiT-NHV (E) and GI-SmBiT (F). The ligation reaction was conducted in PBS with 0.1% BSA, and initial velocity was calculated from the ligation product generated within 5 min according to the measured bioluminescence and the specific activity of the recombinant ligation product. The velocity data for LgBiT-NHV were fitted to the function of V = V_max_[S]/(K_m_+[S]), and those for GI-SmBiT were fitted to the function of V = V_max_[S]^n^/(K_m_^n^+[S]^n^), using SigmaPlot10.0 software. (G) Monitoring the ligation of LgBiT-NHV with GI-SmBiT by bioluminescence and SDS-PAGE. The ligation reaction was conducted in PBS without BSA. At the indicated times, 10 μl of the reaction mixture was removed and either diluted 100,000-fold using the diluting buffer for bioluminescence measurement or mixed with 15 μl of the loading buffer for SDS-PAGE analysis. Thereafter, 10 μl of the diluted sample was used for the bioluminescence measurement or for SDS-PAGE analysis.

Subsequently, we determined the K_m_ values of the circular sortase A towards both substrates. Towards the substrate LgBiT-long-LPETG, the measured initial velocity increased in a linear manner as its concentration increased (Fig. 3E, inner panel), suggesting that the K_m_ value toward the LPETG substrate is too high to be determined in the substrate concentration range used. Although previous fluorescence-based assays suggested that the recombinant sortase A has quite low K_m_ values towards the LPETG substrate, ranging from 16 μM to 140 μM [29,30,31], the HPLC-based assay demonstrated that the low K_m_ values are actually caused by an inner filter effect of the fluorescence assay and the real K_m_ value is as high as ∼5.5 mM [32]. Our present result also suggested that the K_m_ value of sortase A towards the LPETG substrate was quite high.

Toward the 4G-SmBiT substrate, sortase A displayed a sigmoidal saturation curve (Fig. 3E), with an apparent K_m_ value of approximately 1.4 μM and a Hill coefficient of approximately 2.7. The present K_m_ value for 4G-SmBiT was much lower than previously determined values (16–140 μM) for oligoglycine substrates [29,32]. It seemed that the low K_m_ value and the cooperation effect were caused by transient binding of 4G-SmBiT to the other LgBiT substrate.

To confirm the sortase A-catalyzed ligation, we analyzed the ligation product by both bioluminescence and SDS-PAGE (Fig. 2F,G). According to the measured bioluminescence, the concentration of the ligation product reached a plateau (approximately 40 μM) at around 6 h (Fig. 2F), suggesting that almost 80% of LgBiT-long-LPETG was converted to the ligation product. As analyzed by SDS-PAGE (Fig. 2G), the density of the LgBiT-long-LPETG band (indicated by an asterisk) decreased gradually, meanwhile the density of a slightly larger band (presumably the ligation product, indicated by an octothorpe) increased gradually, confirming the ligation of LgBiT-long-LPETG and 4G-SmBiT. Unfortunately, the band of the ligation product overlapped with that of the circular sortase A. In summary, we successfully developed a novel bioluminescent activity assay for sortase A using the recombinant LgBiT-long-LPETG protein and the synthetic 4G-SmBiT tag as a ligation pair.

### 3.2. Validation of the novel bioluminescent activity assay using plant butelase-1

To validate the bioluminescent activity assay using butelase-1, we introduced its recognition motif (NHV) into the C-terminus of LgBiT via a short linker (Fig. 3A). After overexpression and purification, the resultant LgBiT-NHV showed a major band on SDS-PAGE and emitted strong bioluminescence after HiBiT complementation (Fig. 3B), with a specific activity of 5.6 × 10^5^ RLU/fmol, almost identical to that of the native LgBiT. Without HiBiT complementation, its bioluminescence was ∼1000-fold lower (Fig. 3B). Moreover, the expected ligation product, designated as LgBiT-NGI-SmBiT, was almost as active as that of the HiBiT-complemented LgBiT-NHV (Fig. 3B), with a specific activity of 4.9 × 10^5^ RLU/fmol. Thus, the recombinant LgBiT-NHV and the synthetic GI-SmBiT form a suitable ligation pair for developing the bioluminescent activity assay for butelase-1.

Thereafter, we tested whether LgBiT-NHV and GI-SmBiT could be ligated together by the crude extract from tender leaves and stem of butterfly peas (*C. ternatea*). We prepared the crude extract and conducted the ligation reaction using either the reductive buffer reported in previous papers [17,29] or using PBS buffer. After the recombinant LgBiT-NHV, the synthetic GI-SmBiT, and the crude extract were mixed together, a significant increase in bioluminescence was observed, using both the reductive buffer and the PBS buffer (Fig. 3C). In contrast, no bioluminescence increase was observed in the absence of the crude extract or GI-SmBiT in both buffers. Thus, it seemed that butelase-1 in the crude extract can catalyze ligation of the recombinant LgBiT-NHV and the synthetic GI-SmBiT. However, the measured bioluminescence did not increase in a linear manner, especially for the groups with high percentage of the crude extract. This phenomenon was likely caused by proteases in the crude extract that can hydrolyze the ligation product and substrates. In these experiments, no protease inhibitors were added because they might inactivate the peptide ligase. In addition, the crude extract displayed higher ligation activity in PBS than in the reductive buffer. According to specific activity of the expected ligation product (4.9 × 10^5^ RLU/fmol), the maximal concentration of the ligation product was ∼0.1 μM, suggesting that ∼2% of LgBiT-NHV was ligated with GI-SmBiT under catalysis using the crude extract (Fig. 2C).

Subsequently, we conducted the ligation assay using partially purified butelase-1. Under catalysis of the partially purified butelase-1, the concentration of the ligation product increased in an almost linear manner, especially at the initial stage (Fig. 3D). Towards the LgBiT-NHV substrate, butelase-1 displayed a normal hyperbolic saturation curve (Fig. 3E), with a calculated K_m_ value of approximately 27 μM. Towards the GI-SmBiT substrate, butelase-1 displayed a sigmoidal saturation curve (Fig. 3F), with a calculated K_m_ value of approximately 7.3 μM and a Hill coefficient of approximately 2.2. It seemed that the Hill coefficient originated from the binding of GI-SmBiT with LgBiT-NHV during ligation, a phenomenon similar to that observed in the sortase A-catalyzed ligation of LgBiT-long-LPETG and 4G-SmBiT (Fig. 2E).

To confirm that butelase-1 catalyzed the ligation of LgBiT-NHV and GI-SmBiT, we monitored ligation by both bioluminescence and SDS-PAGE (Fig. 3G). According to the measured bioluminescence, the ligation product increased gradually and reached approximately 14 μM at 5 h, suggesting that almost 60% of LgBiT-NHV had been converted to the ligation product. As analyzed by SDS-PAGE (Fig. 3G, inner panel), the density of the LgBiT-NHV band decreased gradually during ligation, meanwhile a slightly larger band (presumably the ligation product, indicated by an asterisk) appeared and its density increased gradually. Thus, it seemed that butelase-1 can catalyze ligation of the recombinant LgBiT-NHV protein and the synthetic GI-SmBiT tag, suggesting that the novel bioluminescent activity assay can accurately monitor the ligation activity of butelase-1.

### 3.3. Screening possible AEP-type peptide ligase from crude extract of some plants

As shown above, the bioluminescent activity assay could detect the ligase activity in a crude extract of butterfly peas sensitively. Thus, we attempted to use this assay to screen novel AEP-type peptide ligase from some other plants. We collected leaves and/or flowers of 80 common higher plants growing around the campus of Tongji University (Table S2). For the possible new peptide ligases in these plants, we knew almost nothing about their recognition motifs and preferred nucleophilic motifs, because neither peptide ligases nor cyclic peptides have been identified from them. Considering that the substrate requirement of peptide ligases is generally not stringent, we used LgBiT-NHV and GI-SmBiT as an initial screening pair that might be recognized by some AEP-type peptide ligases in these plants. In the first round screening, we used 5 μM of LgBiT-NHV and 10 μM of GI-SmBiT as substrates, and 1/2 volume of the crude extract as the enzyme source. If the measured bioluminescence at 5 min or 10 min was significantly higher (over 2-fold) than that at 0 min, the sample was regarded as positive and subjected to further experiments to confirm the presence of the peptide ligase in the crude extract.

In the presence of the crude extract from flowers and flower stalks of *Zephyranthes candida*, an Amaryllidaceae family plant originated in South America and has been introduced to China as a horticultural plant (Fig. 4F), significantly increased bioluminescence was observed (Fig. 4A). In contrast, no bioluminescence increase was observed in the absence of the crude extract or GI-SmBiT (Fig. 4A). Thus, it seemed that the flowers and/or flower stalks of *Z. candida* likely contain an AEP-type peptide ligase that can use LgBiT-NHV and GI-SmBiT as substrates. However, we did not detect significant peptide ligase activity from the crude extract of leaves or corms of *Z. candida*, suggesting that the peptide ligase is primarily expressed in the flowers and/or the flower stalks of this plant.

**Fig. 4.**
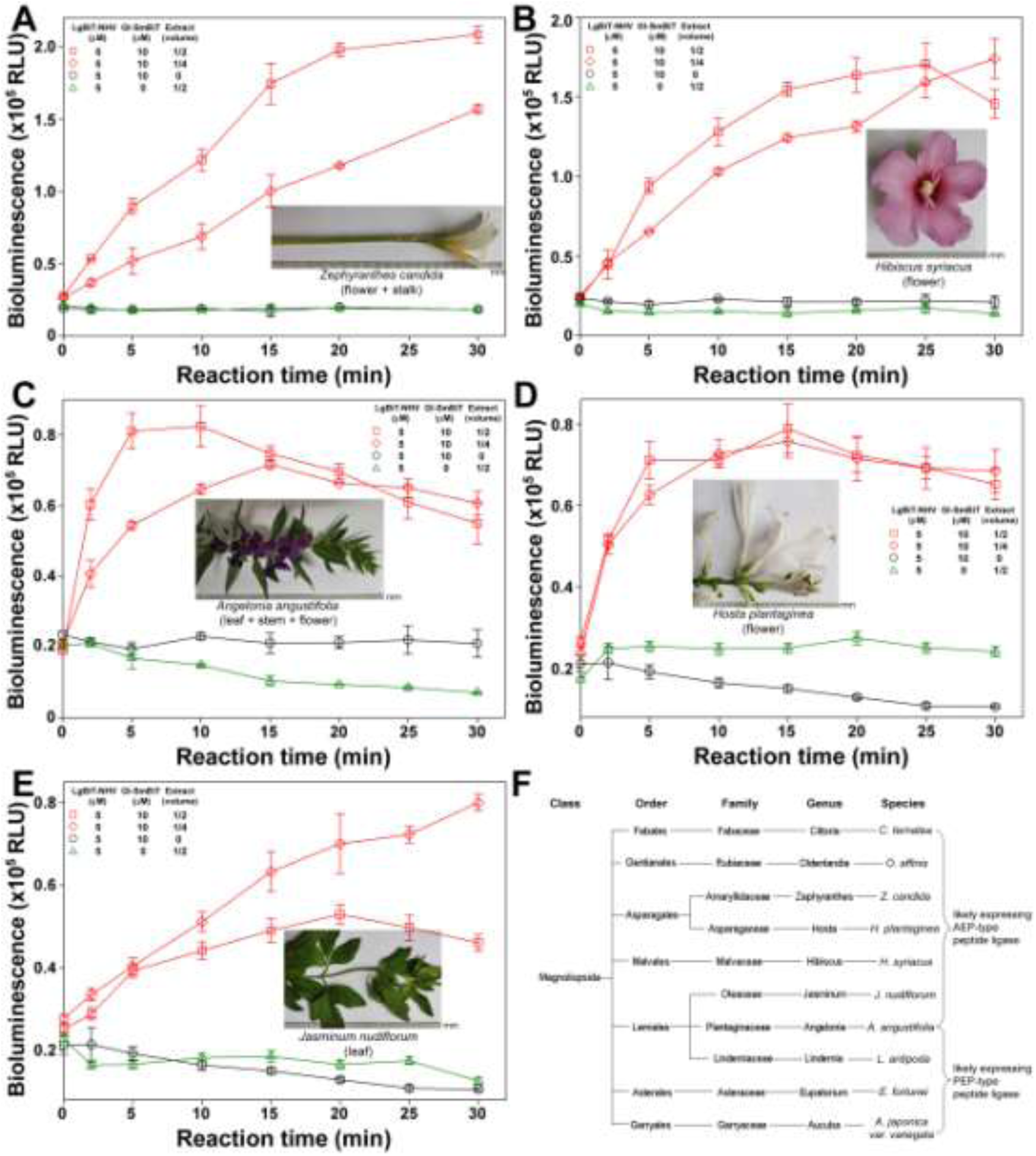
Screening possible AEP-type peptide ligase from crude extracts of some plants using LgBiT-NHV and GI-SmBiT as a ligation pair. (A–E) Ligation activity assays of the crude extracts from *Zephyranthes candida* (A), *Hibiscus syriacus* (B), *Angelonia angustifolia* (C), *Hosta plantaginea* (D), and *Jasminum nudiflorum* (E). Inner panels: Photograph of the plant part used for the activity assay. After the plant part was ground in cold PBS (1 g of plant sample in of 2 ml PBS), the supernatant was used for the ligation assay. At the indicated times, 10 μl of the reaction mixture was removed and diluted 3,000-fold using the diluting buffer. Thereafter, diluted samples were transferred to a 384-well plate (10 μl/well), and bioluminescence was measured immediately on an iD3 microplate reader after addition of the diluted NanoLuc substrate (10 μl/well). The measured bioluminescence data are expressed as the mean ± SD (*n* = 3) and plotted using the SigmaPlot10.0 software. (F) Classification of higher plants likely expressing the AEP-type or PEP-type peptide ligases. *C. ternatea* and *O. affinis* were identified as expressing the AEP-type peptide ligase in previous studies [12,13], while the other plants were identified in the present study.

In the crude extract from flowers of *Hibiscus syriacus*, a Malvaceae family horticultural plant (Fig. 4F), we also detected high peptide ligase activity using LgBiT-NHV and GI-SmBiT as the ligation pair (Fig. 4B). However, the crude extract from leaves of *H. syriacus* had almost no activity, implying that the peptide ligase is mainly expressed in the flowers of *H. syriacus*. The measured peptide ligase activity in the crude extracts of *Z. candida* and *H. syriacus* was similar to that in the crude extract of *C. ternatea* (butterfly pea), implying that catalytic efficiency of their peptide ligase might be as high as that of butelase-1.

In the crude extract from the leaves, stems, and flowers of *Angelonia angustifolia*, a Plantaginaceae family horticultural plant (Fig. 4F), we also detected significant peptide ligase activity (Fig. 4C), although its activity was slightly lower than that in the crude extracts of *Z. candida* and *H. syriacus*. In the crude extract from flowers of *Hosta plantaginea* or from leaves of *Jasminum nudiflorum*, significant peptide ligase activity was also detected (Fig. 4D,E), suggesting that these plants likely express an AEP-type peptide ligase.

From the crude extract of 80 common higher plants, we identified possible AEP-type peptide ligase activity in five evolutionarily distant plants, belonging to three Orders and five Families (Fig. 4F). In future studies, we will purify some of them and confirm whether they are AEP-type peptide ligase or not. In the crude extract of other plants, no significant peptide ligase activity was detected using LgBiT-NHV and GI-SmBiT as a ligation pair, suggesting they might not have an AEP-type peptide ligase. However, we could not exclude the possibility that: 1) Their AEP-type peptide ligase cannot use LgBiT-NHV and GI-SmBiT as substrates; or 2) their AEP-type peptide ligase is only expressed in a special organ or at a special time. Thus, it seemed that the AEP-type peptide ligase is not so rare in higher plants. We estimated that AEP-type peptide ligases might be present in approximately 5–10% of total higher plants, but are likely distributed in evolutionarily distant species (Fig. 4F). Thus, a suitable large-scale screening approach, such as our present bioluminescent activity assay, is needed to identify these AEP-type peptide ligases in future studies.

### 3.4. Screening possible PEP-type peptide ligase from crude extract of some plants

Besides the AEP-type peptide ligase, PEP-type peptide ligase has also been identified from some plants. As shown in Fig. 5A, some cyclic peptides previously identified from *Saponaria vaccaria* and *Pseudostellaria heterophylla* contain a proline residue at the C-terminus of their mature peptide [33,34]. According to their amino acid sequence, we designed a possible ligation pair for screening PEP-type peptide ligase from the crude extract of some plants. A possible recognition motif with the amino acid sequence of Pro-Ile-Gln (PIQ) was introduced into the C-terminus of LgBiT via a short linker, and an aromatic Phe residue was used as a possible nucleophilic motif and introduced to the N-terminus of the synthetic SmBiT via a short linker (Fig. 5B). After overexpression and purification, the resultant LgBiT-PIQ displayed low intrinsic luciferase activity, but its luciferase activity was restored by HiBiT complementation, with a calculated specific activity of approximately 4.3 × 10^5^ RLU/fmol when measured on an iD3 microplate reader using a 384-well plate.

**Fig. 5.**
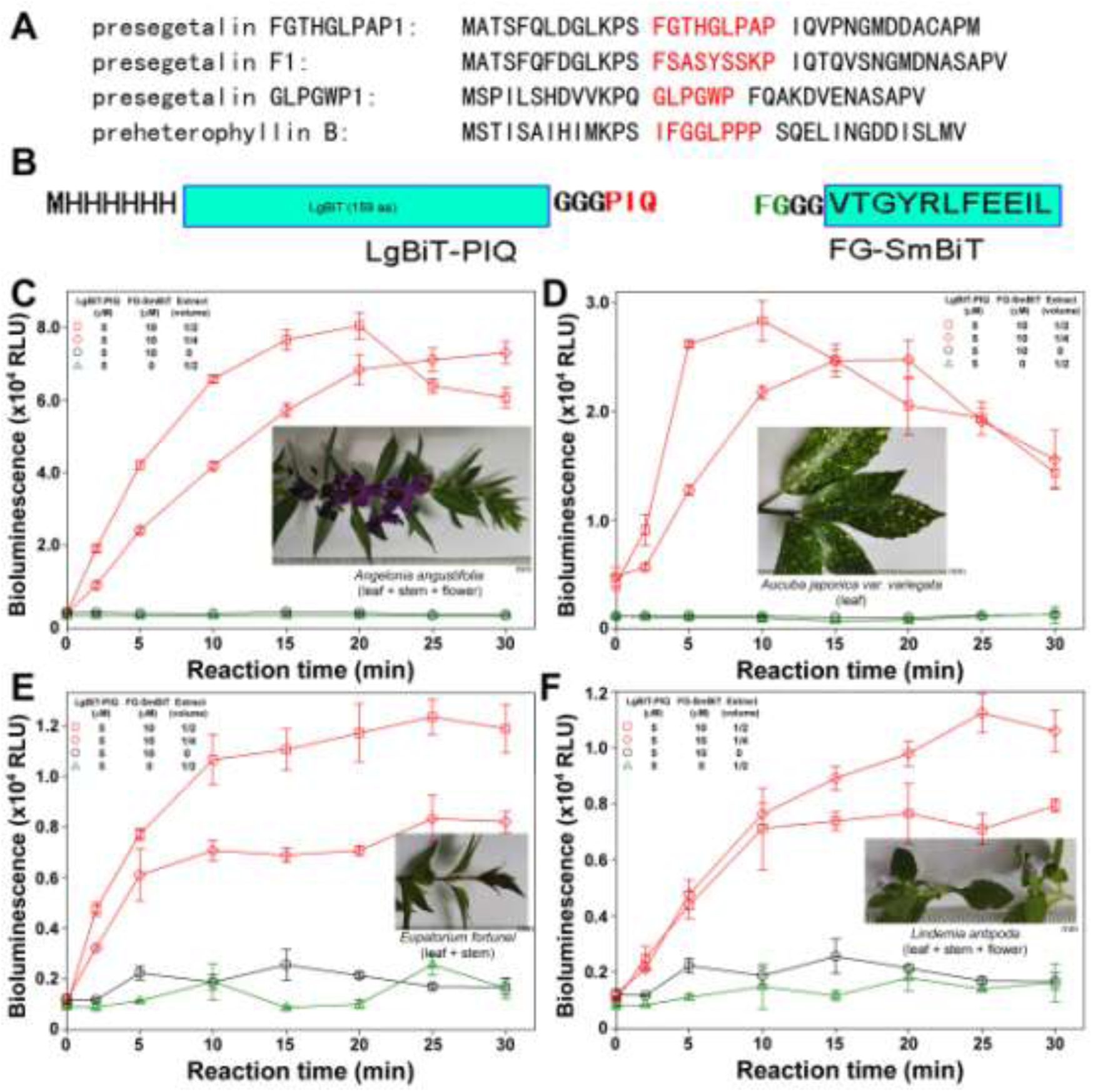
Screening possible PEP-type peptide ligase from the crude extracts of some plants. (A) Amino acid sequence of the precursors of some cyclic peptides carrying a proline residue at the C-terminus of their mature peptide. The mature peptide in the precursors is shown in red. (B) Schematic presentation of the designed ligation pair for screening the PEP-type peptide ligase. (C–F) Ligation activity assay of the crude extracts from *Angelonia angustifolia* (C), *Aucuba japonica* var. *variegata* (D), *Eupatorium fortunei* (E), and *Lindernia antipoda* (F). Inner panels: Photograph of the plant part used for the activity assay. After the plant part was ground in cold PBS (1 g of plant sample in of 2 ml PBS), the supernatant was used for the ligation assay. At the indicated times, 10 μl of the reaction mixture was removed and diluted 2,000-fold using the diluting buffer. Thereafter, diluted samples were transferred to a 384-well plate (10 μl/well), and bioluminescence was measured immediately on an iD3 microplate reader after addition of the diluted NanoLuc substrate (10 μl/well). The measured bioluminescence data are expressed as the mean ± SD (*n* = 3) and plotted using the SigmaPlot10.0 software.

We expected that the bioluminescence would increase once the inactive LgBiT-PIQ protein was ligated with the synthetic low-affinity FG-SmBiT tag by a PEP-type peptide ligase in the crude extract of some plants. After first around screening of the crude extract from 80 common higher plants (Table S2), we confirmed that four of them likely expressed PEP-type ligase activity (Fig. 1C–F). After LgBiT-PIQ, FG-SmBiT, and the crude extract from *Angelonia angustifolia* were mixed together, a marked increase in bioluminescence was observed, but no bioluminescence increase was observed in the absence of the crude extract or FG-SmBiT (Fig. 4C). Thus, it seemed that the crude extract of this plant likely contains a PEP-type peptide ligase that can catalyze ligation of LgBiT-PIQ and FG-SmBiT. In the crude extract from *Aucuba japonica var. variegate*, bioluminescence increase was also observed (Fig. 1D), suggesting this plant might also express a PEP-type peptide ligase. In the crude extract from *Eupatorium fortunei* and *Lindernia antipoda*, we also detected possible PEP-type peptide ligase activity (Fig. 1E,F), although their activities were lower than that of the crude extract from *Angelonia angustifolia* and *Aucuba japonica var. variegate*.

By designing a possible ligation pair based on the amino acid sequence of certain cyclic peptides and their linear precursors, we identified possible PEP-type peptide ligase activity in the crude extract of four evolutionarily distant plants that belong to three Orders and four Families (Fig. 4F). Thus, it seemed that the PEP-type peptide ligase is also not so rare in higher plants, and more of them will be discovered in future studies via large-scale screening using the novel bioluminescent activity assay.

## 4. Discussion

In the present study, we developed a novel bioluminescent activity assay to characterize known peptide ligases or to screen for new ones from crude extracts of various plants or other sources. To set up the bioluminescent activity assay, a recognition motif of one or a group of peptide ligase(s) should be genetically fused to the C-terminus of the inactive LgBiT protein, and a nucleophilic motif preferred by the peptide ligase(s) should be chemically introduced into the N-terminus of the synthetic SmBiT tag. For known peptide ligases, their recognition motifs and preferred nucleophilic motifs are largely known. For new peptide ligases, their recognition motifs and preferred nucleophilic motifs can be deduced from their products and substrates, such as the mature cyclic peptides and their linear precursors. A short sequence at the C-terminal conjunction of the mature peptide and its precursor can be used as the recognition motif, and several amino acids at the N-terminus of the mature peptide can be used as the nucleophilic motif. Considering that peptide ligases typically have a wide substrate spectrum, the designed ligation version LgBiT and SmBiT would be used for characterizing or screening a group of peptide ligase. Once the ligation version LgBiT proteins were overexpressed and purified, their intrinsic luciferase activity should be low, but their activity after HiBiT complementation should be high. If so, the recombinant LgBiT proteins are suitable for developing a peptide ligation assay.

Compared with previously reported peptide ligase activity assays, such as the HPLC-based, fluorescence-based, and mass spectrometry-based assays, the present bioluminescent activity assay has several advantages. (1) The present activity assay is ultrasensitive, because the ligation product restores high luciferase activity and can be quantified accurately according to its bioluminescence, with a detection limit of femtomoles, or even lower, on most microplate readers. (2) The present assay is convenient for use because it contains only three major steps (ligation, dilution, and bioluminescence measurement) and typically can be finished within one hour. Moreover, this assay can deal with large numbers of samples, and thus is especially suitable for large-scale screening of novel peptide ligases from various plants or other sources. This assay does not require expensive equipments, and is thus suitable for most laboratories. (3) This assay is resistant to interference of proteases, and thus is especially suitable for screening novel peptide ligases from crude extracts. Proteases are widely present in crude extracts and seriously interfere with screening via hydrolysis of the ligation products and substrates. However, protease inhibitors typically cannot be used in screening because peptide ligases are a special type of proteases, and thus might be inactivated by protease inhibitors. For the bioluminescent activity assay, hydrolysis of LgBiT and SmBiT had no serious effects, because only the ligation product can emit bioluminescence. Once LgBiT and SmBiT were ligated together, the ligation product would become more stable and more resistant to protease hydrolysis.

Peptide ligases have a wide range of applications for protein labeling or cyclic peptide synthesis. However, to date, only a few peptide ligases have been identified, mainly because of the lack of an efficient and convenient assay for large-scale screening. Previous studies typically focused on a few plants that produce cyclic peptides. Unfortunately, the identification of cyclic peptides is also difficult and time-consuming. Thus, it remains a challenge to quickly test whether a plant expresses a peptide ligase or not. Fortunately, the present study provides a practical and efficient approach to solve this problem. Our activity assay only needs 0.1–1.0 g of plant sample (flowers, leaves, or other parts), and typically, a single researcher can deal with hundreds of samples within 1–2 weeks. Thus, the bioluminescent activity assay is especially suitable for large-scale screening. By screening the crude extract from 80 common higher plants, we identified that five of them likely express AEP-type peptide ligase and four of them likely express PEP-type peptide ligase (Fig. 4F). Thus, it appeared that peptide ligases are not so rare in higher plants, and more of them are waiting to be discovered in the future. In addition to screening, the bioluminescent activity assay would also play a critical role in subsequent purification and characterization of these novel peptide ligases in our future studies, because it can follow these peptide ligases during the purification steps according to their activity. Besides the AEP-type and PEP-type peptide ligases, other types of peptide ligases might be also present, because cyclic peptides have highly divergent amino acid sequences [33,35,36]. By designing suitable LgBiT and SmBiT ligation pairs, more peptide ligases with difference substrate preference could been screened and identified in future studies. In addition, novel cyclic peptides likely can be identified from those plants expressing peptide ligases. Thus, our present bioluminescent activity assay opens the door for the rapid discovery of novel peptide ligases or even novel cyclic peptides from various plants or other sources.

## Abbreviations

4G-SmBiT: a ligation version SmBiT tag carrying four glycine residues at its N-terminus
AEP: asparaginyl endopeptidase
BSA: bovine serum albumin
EDTA: ethylene diamine tetraacetic acid
FG-SmBiT: a ligation version SmBiT tag carrying the sequence of Phe-Gly-Gly-Gly at its N-terminus
GI-SmBiT: a ligation version SmBiT tag carrying the sequence of Gly-Ile-Gly-Gly at its N-terminus
HiBiT: a high-affinity complementation tag for NanoBiT
HPLC: high performance liquid chromatography
LgBiT: A large NanoLuc fragment for NanoBiT
LgBiT-long-LPETG: a ligation version LgBiT protein carrying the sequence of Leu-Pro-Glu-Thr-Gly-Gly at its C-terminus via a long linker with 18 residues
LgBiT-LPETG: a ligation version LgBiT protein carrying the sequence of Leu-Pro-Glu-Thr-Gly-Gly at its C-terminus via a linker of three Gly residues
LgBiT-NHV: a ligation version LgBiT protein carrying the sequence of Asn-His-Val at its C-terminus via a linker of three Gly residues
LgBiT-PIQ: a ligation version LgBiT protein carrying the sequence of Pro-Ile-Gln at its C-terminus via a linker of three Gly residues
NanoBiT: NanoLuc Binary Technology
PAL: peptidyl asparaginyl ligase
PBS: phosphate-buffered saline
PCR: polymerase chain reaction
PEP: Prolyl endopeptidase
RLU: relative luminescence unit
SDS-PAGE: sodium dodecyl sulfate-polyacrylamide gel electrophoresis
SmBiT: a low-affinity complementation tag for NanoBiT
UV: ultra-violet.

## Acknowledgments

This work was supported by grant from the National Natural Science Foundation of China (31971193).

## Supplementary materials

**Table S1:**
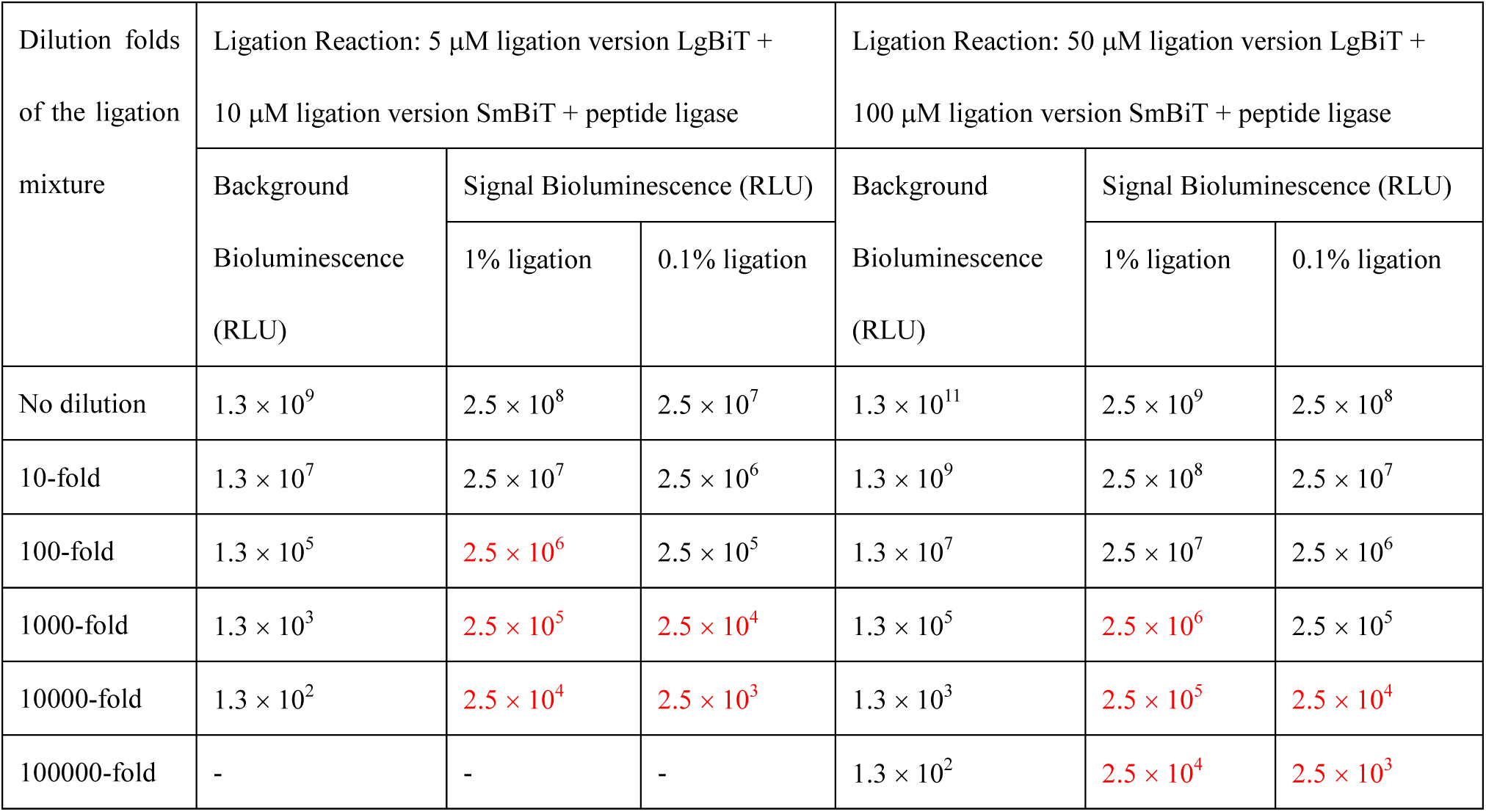
The calculated theoretical values of the background bioluminescence originated from the SmBiT-complemented LgBiT and the signal bioluminescence originated from the ligation product. Presuming that K_d_ value of the ligation version LgBiT with the ligation version SmBiT is identical to that of the original LgBiT–SmBiT pair (K_d_ = 190 μM). Presuming that 1% or 0.1% of the ligation version LgBiT was ligated with the ligation version SmBiT by the peptide ligase. Presuming that the specific luciferase activity of both the ligation product and the SmBiT-complemented LgBiT was 5×10^5^ RLU/fmol when measured on an iD3 microplate reader using a 384-well plate. After the ligation, the reaction mixture was diluted as indicated as in the table and the diluted samples were transferred to a 384-well plate (10 μl/well). After addition of the diluted NanoLuc substrate (10 μl/well), bioluminescence was immediately measured on an iD3 microplate reader under the luminescence mode. Those signal data significantly higher than the corresponding background data are shown as red.

**Table S2:** Summary of the 80 higher plants used for screening the AEP-type peptide ligase and PEP-type peptide ligase in the present study.

| Plant name | Extraction parts | Possible AEP-type peptide ligase activity | Possible PEP-type peptide ligase activity |
| --- | --- | --- | --- |
| *Cercis chinensis* | leaves | - | - |
| *Campsis grandiflora* | leaves | - | - |
| *Salvia japonica* | leaves + flowers | - | - |
| *Lycoris radiata* | flowers | - | - |
| *Solanum nigrum* | leaves | - | - |
| *Loropetalum chinense* var. *rubrum* | leaves | - | - |
| *Erigeron annuus* | leaves | - | - |
| *Cayratia japonica* | leaves | - | - |
| *Oxalis triangularis* subsp. *papilionacea* | leaves | - | - |
| *Lactuca indica* | leaves | - | - |
| *Euphorbia humifusa* | leaves | - | - |
| *Setaria viridis* | leaves | - | - |
| *Calystegia hederacea* | leaves | - | - |
| *Portulaca umbraticola* | leaves + flowers | - | - |
| *Lagerstroemia indica* | leaves + flowers | - | - |
| *Commelina communis* | leaves | - | - |
| *Echinochloa crusgalli* | leaves | - | - |
| *Eriobotrya japonica* | leaves | - | - |
| *Parthenocissus tricuspidata* | leaves | - | - |
| *Aucuba japonica* var. *variegata* | leaves | - | + |
| *Hedera helix* | leaves | - | - |
| *Eupatorium fortunei* | leaves | - | + |
| *Agapanthus africanus* | leaves | - | - |
| *Canna indica* | leaves + flowers | - | - |
| *Begonia cucullata* var. *hookeri* | leaves + flowers | - | - |
| *Phyllanthus urinaria* | leaves | - | - |
| *Pinellia ternata* | leaves | - | - |
| *Morus alba* | leaves | - | - |
| *Hibiscus syriacus* | flowers | + | - |
| *Hibiscus syriacus* | leaves | - | - |
| *Potentilla supina* | leaves | - | - |
| *Metaplexis japonica* | leaves | - | - |
| *Stephania japonica* | leaves | - | - |
| *Ulmus pumila* | leaves | - | - |
| *Vitex negundo* | leaves | - | - |
| *Sedum emarginatum* | leaves | - | - |
| *Portulaca grandiflora* | leaves | - | - |
| *Lantana camara* | leaves | - | - |
| *Plumbago auriculata* | leaves | - | - |
| *Gaura lindheimeri* | leaves | - | - |
| *Mucuna sempervirens* | leaves | - | - |
| *Wisteria sinensis* | leaves | - | - |
| *Salvia leucantha* | leaves | - | - |
| *Mirabilis jalapa* | leaves + flowers | - | - |
| *Zephyranthes candida* | flowers + stalks | + | - |
| *Zephyranthes candida* | leaves | - | - |
| *Sonchus wightianus* | leaves | - | - |
| *Pterocarya stenoptera* | leaves | - | - |
| *Trifolium repens* | leaves + flowers | - | - |
| *Taxus wallichiana* var. *chinensis* | leaves | - | - |
| *Gardenia jasminoides* | leaves | - | - |
| *Farfugium japonicum* | leaves | - | - |
| *Elephantopus scaber* | leaves | - | - |
| *Oxalis corniculata* | leaves | - | - |
| *Viburnum odoratissimum* | fruits | - | - |
| *Impatiens walleriana* | leaves | - | - |
| *Acalypha australis* | leaves | - | - |
| *Alternanthera philoxeroides* | leaves | - | - |
| *Tulbaghia violacea* | flowers + stalks | - | - |
| *Hydrangea macrophylla* | leaves | - | - |
| *Euonymus fortunei* | leaves | - | - |
| *Aster ageratoides* | leaves | - | - |
| *Ajania pacifica* | leaves | - | - |
| *Hosta plantaginea* | flowers | + | - |
| *Robinia pseudoacacia* | leaves | - | - |
| *Phytolacca acinosa* | leaves | - | - |
| *Broussonetia papyrifera* | leaves | - | - |
| *Kyllinga brevifolia* | leaves | - | - |
| *Acer palmatum* | leaves | - | - |
| *Angelonia angustifolia* | leaves + stems + flowers | + | + |
| *Pentas lanceolata* | leaves + flowers | - | - |
| *Torenia fournieri* | leaves | - | - |
| *Lindernia antipoda* | leaves + flowers | - | + |
| *Jasminum nudiflorum* | leaves | + | - |
| *Zamioculcas zamiifolia* | leaves | - | - |
| *Lobelia sessilifolia* | leaves | - | - |
| *Zephyranthes carinata* | flowers + stalks | - | - |
| *Clerodendrum trichotomum* | leaves | - | - |
| *Vigna unguiculata* | Pods | - | - |
| *Phaseolus vulgaris* | Pods | - | - |
| *Pisum sativum* | Pods | - | - |
| *Glycine max* | Pods | - | - |

**Fig. S1.**
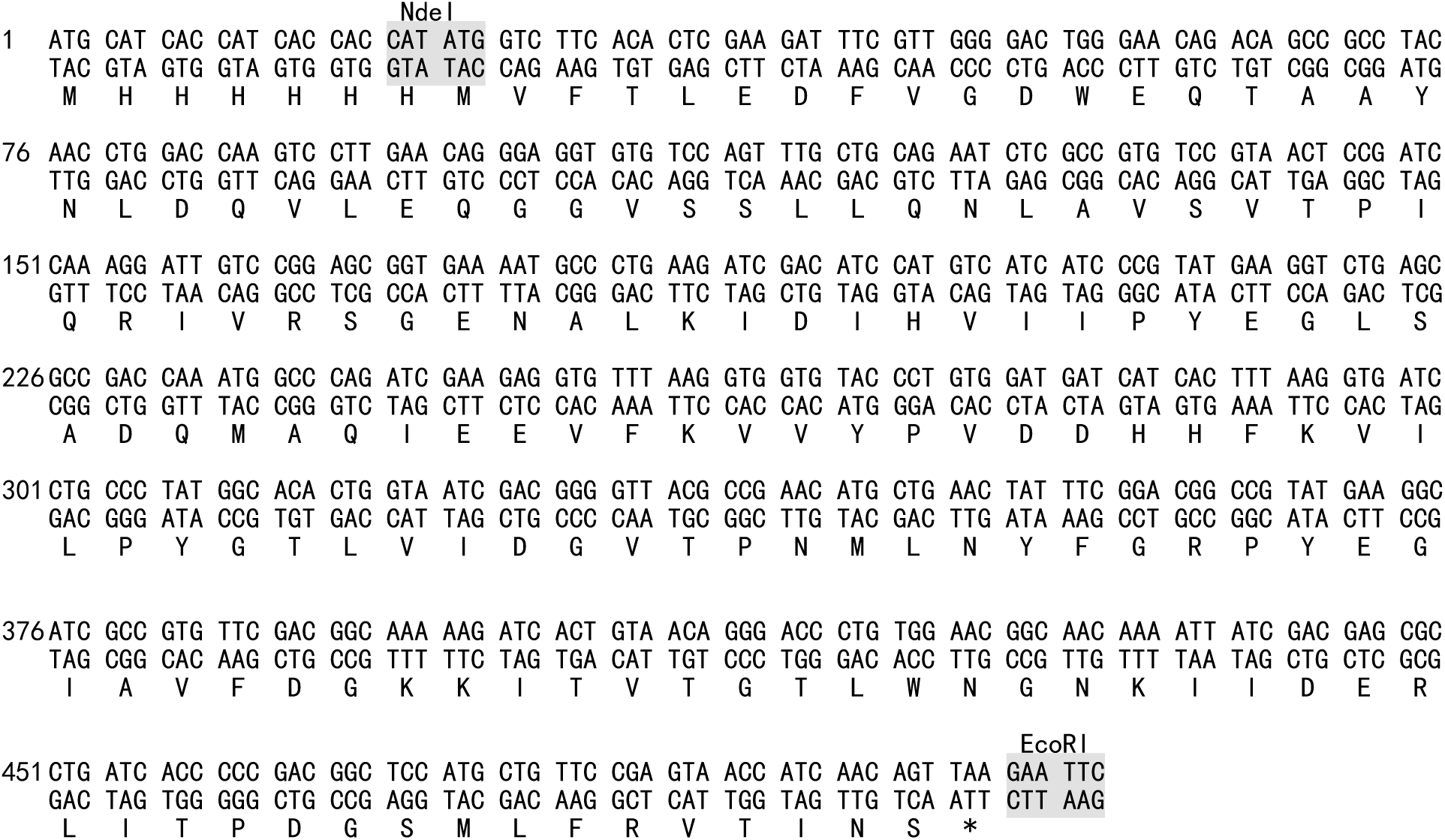
The nucleotide and amino acid sequence of the N-terminally 6×His-tagged original LgBiT protein overexpressed in *E. coli*. The synthetic DNA fragment was cleaved with restriction enzymes NdeI and EcoRI and then ligated into a pET expression vector for recombinant expression of the LgBiT protein.

